# itsfm, an open-source package to reliably segment and measure sounds by frequency modulation

**DOI:** 10.1101/2021.01.09.426033

**Authors:** Thejasvi Beleyur

## Abstract

Analysing animal vocalisations in detail provides insights into the biomechanics, decision making and sensory processes behind their behaviours. Echolocating bats, and in particular, the CF-FM calls of high-duty cycle bats serve as a convenient model system to illustrate this point. The CF component in the CF-FM call is used for prey detection and the FM component is used in target ranging. According to the behavioural context at hand such as flight with conspecifics or prey capture, bats choose to increase the duration, intensity or spectral range of the components differently. Studying the call component alterations requires an objective methodology that first segments the components and then allows measurements on them. Studies till now have segmented the call components manually, or automatically using what I term the ‘peak-frequency’ method. Manual segmentation is error prone, while the ‘peak-frequency’ method requires on-axis recordings for good results. Despite multiple papers using a peak-frequency based segmentation, there remain no publicly available software implementations. `itsfm` is an open-source package that fills this gap with two implemntations that can segment CF-FM calls, one of them being an implementation of the peak-percentage method. `itsfm` additionally introduces the ‘pseudo-Wigner-Ville distribution’ (PWVD) method for call segmentation, thus allowing the segmentation of calls captured under a wider variety of recording conditions. I create a synthetic dataset and assess the performance of the PWVD method and the ‘peak-frequency’ method. The PWVD performs consistently well in call component segmentation in comparison to the peak-percentage method. I also discuss the supporting methods in the `itsfm` package that can help the further automatic segmentation, measurement and analysis of sounds. Though originally developed for the segmentation and measurement of CF-FM bat calls, the methods in `itsfm` are speciesagnostic, and may be used for vocalisations of any type.

## 0.1 Introduction

Vocalisations are a window into the sensory, behavioural and biomechanical states of an animal (Green and Marler 1979; Metzner and Müller 2016). Echolocating bats present a unique model system where vocalisations play a fundamental role in the animal’s sensorimotor decisions. Echolocating bats emit loud calls and listen for returning echoes to detect objects around them (Griffin 1958). Bats are known to flexibly alter various aspects of their calls to optimise echo detection, and thus their own sensory input. For instance, bats flying in the open emit long calls with a narrow bandwidth, and switch to short high-bandwidth sweeps as they are about attack an insect prey (Fenton 2013).

Among echolocating bats, the so-called CF-FM bats are a particularly interesting model system to study sensorimotor decisions. CF-FM calls (Figure 1) consist of a constant-frequency (CF) component and upto two frequency-modulated (FM) components. The CF component is used in the detection of prey wing-flutter (Schnitzler and Denzinger 2011) while the FM component is used in target ranging (Tian and Schnitzler 1997). Bats are known to independently alter the CF and FM components depending on the presence of echolocating conspecifics (Fawcett et al. 2015), artificial playbacks (Lu, Zhang, and Luo 2020; Hage et al. 2013, 2014) or during flight manuevers (Tian and Schnitzler 1997; Schoeppler, Schnitzler, and Denzinger 2018). Studying how CF-FM bats alter their call components requires an objective method that can reliably segment the components, and thus facilitate accurate acoustic parameter measurement.

**Figure 1:**
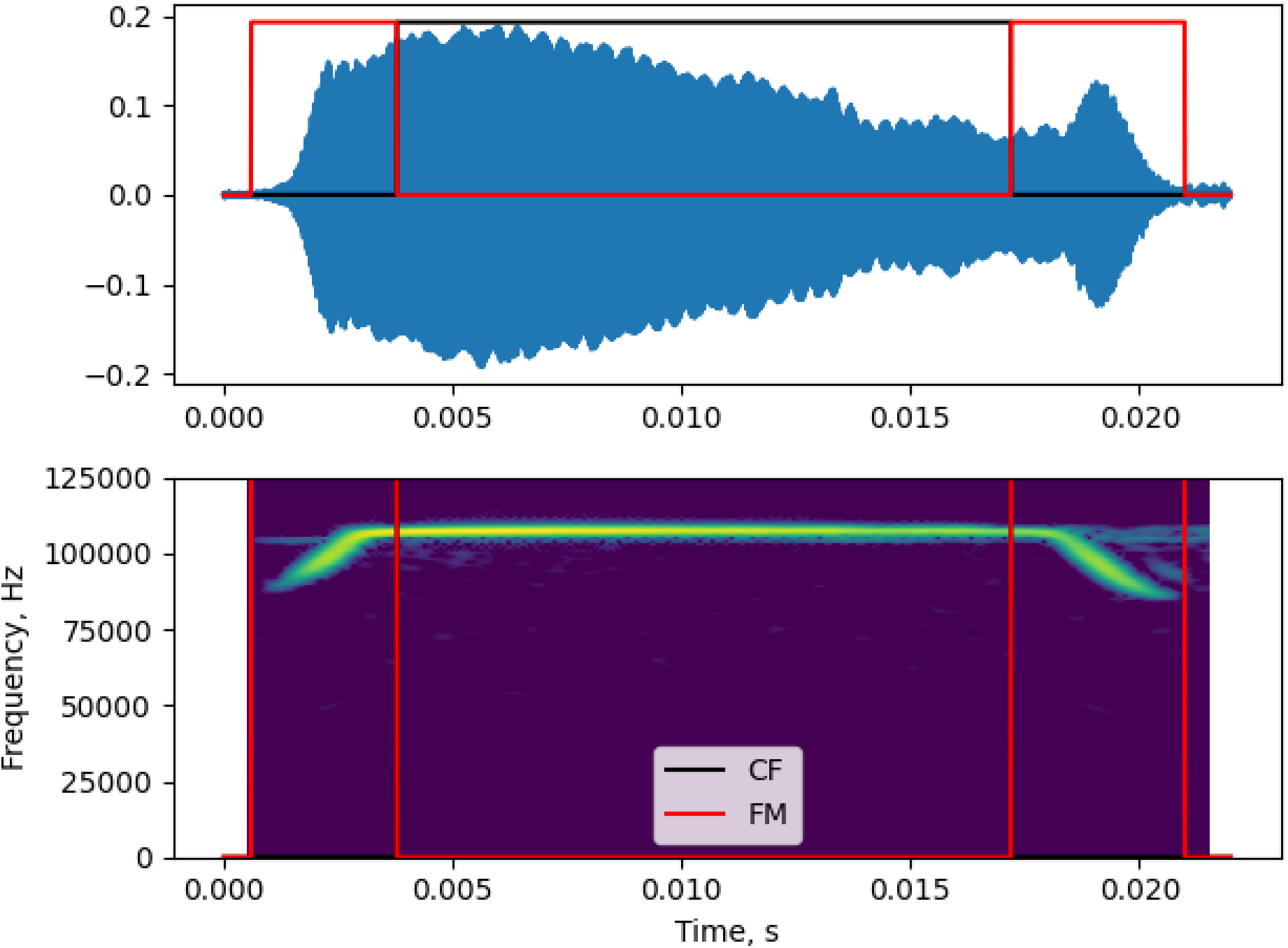
Diagnostic plot showing the CF/FM segmentation output of a *Rhinolophus euryale/mehelyi* call

### 0.1.1 State of the art: CF-FM call segmentation

Manual segmentation of calls into CF and FM is the most intuitive and direct one can take, and has been an approach used in publications to date (Vater et al. 2003; Fawcett et al. 2015; Gessinger et al. 2019). Manual segmentation however doesn’t scale with sample size, are not very reproducible and can be biased (Brumm et al. 2017). Tian and Schnitzler (1997) is to my knowledge, the first publication to attempt a semi-manual segmentation of the CF and FM call components, and their method has formed the founding basis for for further work. I hereby refer to methods based on their approach as the ‘peak percentage’ approach. Tian and Schnitzler (1997) method relies on the fact that the CF component is at the highest frequency and forms a large part of the call. By filtering below and above a threshold frequency close to the CF frequency, the FM and CF components can be separated. In *Rhinolophus ferrumequinum*, Tian and Schnitzler (1997) define the threshold frequency at 0.8kHz below the 2nd harmonic of ‘CF component’. 0.8 kHz corresponds to around 1% of the CF peak frequency (~80 kHz), and is also equivalent to filtering at 99 % of the CF peak frequency. In their semi-manual method Tian and Schnitzler (1997) measured the CF peak frequency using an FFT frequency analyser to separate FM and CF components. Schoeppler, Schnitzler, and Denzinger (2018) further automate the method of Tian and Schnitzler (1997) by using 99% of the CF frequency to define FM components in a spectrogram based method run on a computer. Lu, Zhang, and Luo (2020) follow on the methodology of Schoeppler, Schnitzler, and Denzinger (2018) and Tian and Schnitzler (1997), and set the FM to begin at 97% of the CF peak frequency. Peak-percentage type approaches allow a straightforward segmentation and measurement, however the method was developed keeping on-axis, high signal-to-noise ratio recordings in mind, such as those that are obtained in flight room experiments. For instance, a pre-requisite for the peak-percentage method to work is a spectrally dominant CF component, in the absence of which the threshold frequency is not identified correctly, leading to poor segmentation. The peak-percentage method also requires setting a reasonable peak-percentage to define the threshold frequency that determines where the CF ends and FM begins. Previous studies have used percentages between 97-99% of the CF peak frequency, and the exact percentage is likely to play a big role in segmentation accuracy.

Despite the apparent popularity of the peak-percentage method there are no openly available code implementations that have been tested for their performance against synthetic data. While code descriptions help explaining the principles behind design, it is not sufficient to ensure uniformity or correctness in implementation. Differences in implementation may lead to differences in scientific results (Baker and Vincent 2019; McFee et al. 2018). Publishing code as publicly available packages allows for external code inspection and improvements. `itsfm` fills the gap by implementing the peak-percentage method and introducing an alternate segmenation method. The segmentation methods are tested against synthetic datasets, with a open-source code base written in a non-proprietary language, and supported by a detailed user-guide online.

## 0.2 Package description

`itsfm` currently provides two main approaches to segment the CF and FM components of a sound (Figure 1), the ‘peak-percentage’ and ‘pwvd’ methods.

### 0.2.1 Peak-percentage segmentation

The `peak-percentage` method is best for sounds with one or more dominant CF components of the same frequency, and FM components that are below the CF component’s frequency (Figure 2). A typical rhinolophid/hipposiderid CF-FM call is the simplest example for which this method works. This method’s implementation is inspired by previously published efforts to segment CF-FM calls into their respective components (Lu, Zhang, and Luo 2020; Tian and Schnitzler 1997; Schoeppler, Schnitzler, and Denzinger 2018). The approach implemented here creates two versions of the raw audio that are low and high passed at a threshold frequency. The threshold frequency is calculated as a fixed percentage of the raw audio’s peak frequency, eg. 99%. The dB rms profile of the low and high passed audio are then calculated and compared by subtraction. Continuous regions where the low-passed audio is greater than the high-passed audio are considered FM regions, and CF regions where it is vice-versa (Figure 2).

**Figure 2:**
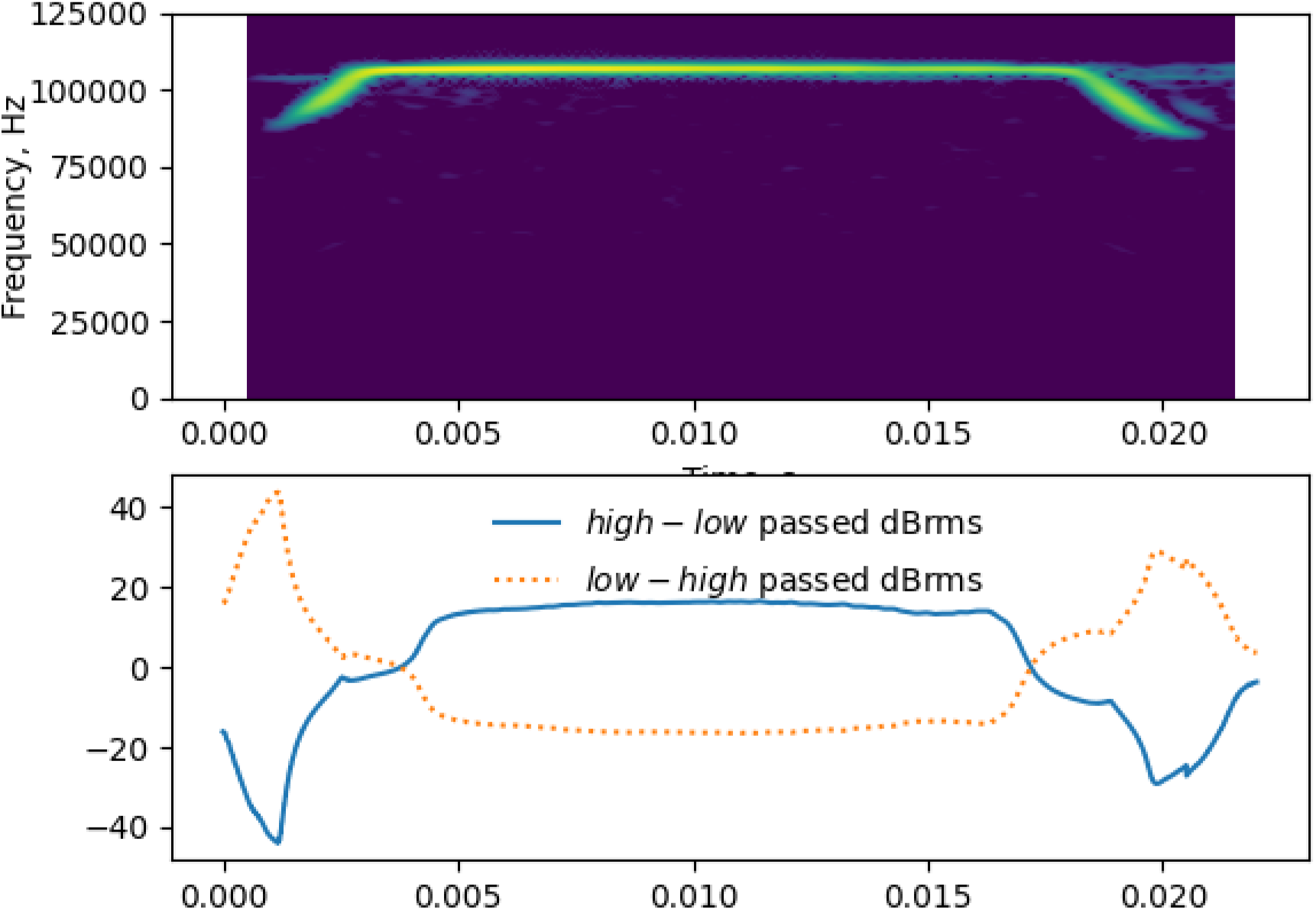
Diagnostic output showing the underlying basis of the peak-percentage CF-FM segmentation method. Top: A spectrogram representation of the call shown in Figure 1, Below: the high/low-passed dB rms profiles of the call. The peak frequency of the entire call is taken and a high and low-passed version of the sound if created at 99% of the call peak frequency. The dB rms profile differences of the high and low-passed sounds are calculated and subtracted from each other. The region where the low-passed dB rms profile is higher are labelled FM and vice-versa as CF.

The peak-percentage method is relatively easy to parameterise as it accepts two intuitive input parameters, the `peak_percentage` (peak percentage value between 0-1) and `window_size` (the number of samples for the window used to calculate the dB rms profile of the high/low passed audio). A set of additional optional parameters may also be specified. The default low/high pass filter is a second order elliptic filter with 3dB ripple (pass band) and 10dB minimum attenuation in the stop band. The user may also optionally specify their own recursive filter coefficients.

A major drawback in the peak-percentage method is its limited use-cases. Sounds must be sufficiently similar to the ‘ideal’ spectro-temporal shape of a classic CF-FM call, or they will be mis-segmented. Not even all CF-FM calls are likely to be segmented properly, eg. CF-FM calls emitted during landing or approach with short CF segments and longer FM segments. If the CF segment of the input sound does not contribute majorly to the spectrum, then the peak-percentage method fails. Experience with field recordings having off-axis CF-FM bat calls shows that the peak-percentage method also fails here because the CF component may not be as dominant as in on-axis recordings of the same call. Aside from CF-FM echolocation calls, the peak-percentage method may also be used for certain types of bird calls with long CF and short FM calls (eg. those emitted by the *Pachycephala* genus)

### 0.2.2 PWVD segmentation

The `pwvd` method (Figure 3) tracks the frequency modulation over the course of the input sound. Regions with an above threshold frequency modulation are considered FM regions, and those below are considered CF regions. The frequency modulation over the course of a sound is estimated by first generating a a sample-level ‘frequency profile’ through the use of the Pseudo Wigner-Ville Distribution (PWVD). The PWVD is a relatively underutilised method in bioacoustics (but see Fu and Kloepper 2018; Kopsinis et al. 2010) which generates time-frequency representations with high spectro-temporal resolution (Boashash 2015). The first derivative of the frequency profile is used to generate a sample-level estimate of frequency modulation and thus segment regions that are above or below the threshold.

**Figure 3:**
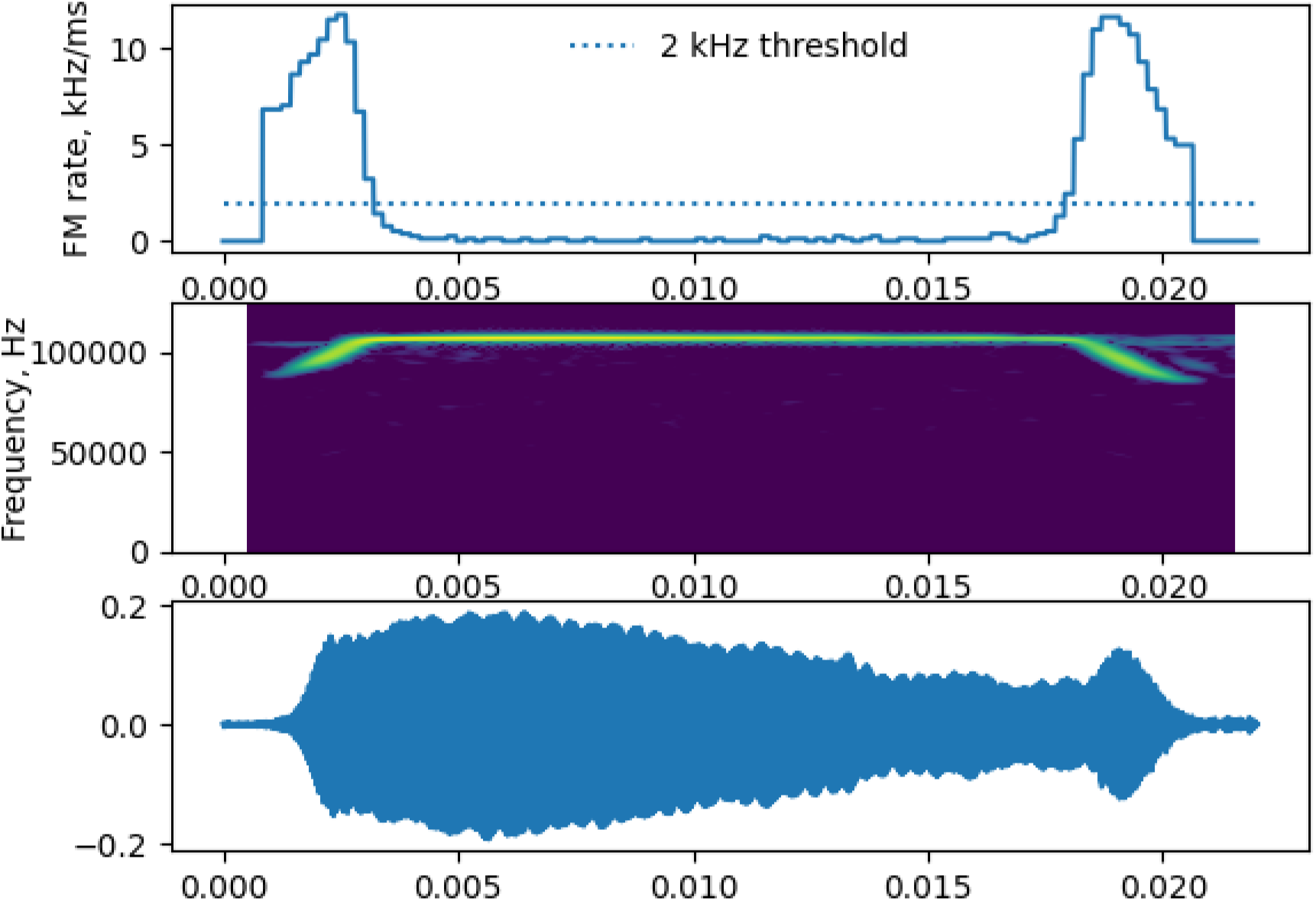
Diagnostic plots of the pwvd method. Top: Sample-level frequency modulatation rate estimates. All regions ≥ the threshold FM rate (here 2kHz/ms) are considered FM regions, while all regions below this are considered CF regions. Middle-Bottom: spectrogram and waveform of the original sound for comparison.

The `pwvd` method requires somewhat more parametrisation and methodological understanding than the `peak-percentage` method. The pwvd method’s effectiveness is dependent on the `fmrate_threshold` (frequency modulation threshold, in kHz/ms), `pwvd_window` size (number of samples used to form the ‘slices’ of the time-frequency representation) and `tfr_cliprange` (permitted dynamic range in dB, used to clip the time-frequency representation and remove noise). In addition to these primary parameters, the pwvd method can be further fine-tuned to improve segmentation. The frequency profile is currently generated by tracking the dominant frequency over each slice of the PWVD representation. The dominant frequency approach is susceptible to noise and changes in sound levels over time, and thus requires additional correction routines that interpolate between problematically tracked regions. The problematic regions are identified by measuring the accelaration (second derivative) of the sound’s frequency profile. Regions above a user-set threshold are considered ‘spiky’ and are interpolated or extrapolated based on neighbouring regions frequency estimates.

Even though the `pwvd` method requires some initial effort to parameterise, the flexibility it provides allows the analysis of a much wider-range of sounds than the `peak-percentage` method. The CF/FM segmentation in the `pwvd` method is independent of the actual call shape, and even complex sounds such as bat social calls and bird songs could be segmented through this method. A major drawback of the current `pwvd` implementation is its inability to reliably segment multi-harmonic sounds. Multi-harmonic sounds present a challenge for the simple frequency tracking in place currently, and alternative algorithms will be a focus of future development.

### 0.2.3 Supporting itsfm methods

Along with the primary segmentation methods, `itsfm` has a collection of supporting methods that allow quantification, visualisation and batch-processing. A series of inbuilt measurement functions allow acoustically relevant measurements such as duration, rms, peak-frequency, or terminal frequency. Custom measurements may also be specified by the user. A sound analysed with the `pwvd` method generates more than the identified CF/FM regions. Raw data on the frequency profile of the sound and the rate of frequency modulation over time are of interest to researchers studying the speed at which vocalisations can be modulated from a behavioural and biomechanical viewpoint (Metzner and Müller 2016; Hage et al. 2013). Along with the background data used to form the segmentations, `itsfm` also provides a series of inbuilt visualisation functions to visualise the input sound itself (`visualise_sound`) and generate diagnostic plots of the segmentation output through the `itsfmInspector` class and `visualise_cffm_segmentation` (Figure 1).

Handling audio recordings made in the field calls for the individual handling of each recording. To aid the reproducible processing of multiple files with unique input parameters `itsfm` can also be called through a command-line interface that accepts batch files in the CSV format. To facilitate iterative parameter optimisation, the user can choose to select only a few audio recordings or the entire set of files defined in the batch file. For each processed audio file, the diagnostic plot and measurements are saved in the working directory.

The `itsfm` package also comes bundled with a series of field recordings of bat calls of various hipposiderid, rhinolophid and noctilionid species. These field recordings allow the user to test the utility of the methods in the package, and gain familiarity with setting correct parameters.

## 0.3 Methods evaluation

### 0.3.1 Synthetic dataset creation and segmentation

To test the accuracy of the segmentation methods implemented in the `itsfm` package, I generated a set of synthetic CF-FM calls with known segment durations and spectral properties. Synthetic calls were generated based on calls broadly based on the structure of rhinolophid and hipposiderid call parameters using the package’s inbuilt `make_cffm_call function`. A set of 324 synthetic calls were made through a combination of parameters in Table 1. Each synthetic call consisted of an iFM, CF and tFM component (naming as per (Tian and Schnitzler 1997)), and is Tukey windowed without any padded silent samples or background noise. All synthetic calls were generated at a sampling rate of 250kHz.

**Table 1:** Parameter values used to generate synthetic CF-FM calls. The parameters broadly reflect the call shape of a rhinolophid/hipposiderid CF-FM bat calls. iFM and tFM regions were generated from the same FM parameter set. 9 CF × 6 iFM × 6 tFM combinations = 324 calls

| Parameter name | Parameter values |  |  |
| --- | --- | --- | --- |
| CF duration (ms) | 5 | 10 | 15 |
| CF peak frequency (kHz) | 40 | 60 | 90 |
| i/t FM duration (ms) | 1 | 2 |  |
| i/t FM bandwidth (kHz) | 5 | 10 | 20 |

The synthetic calls were segmented according to method-specific parameters that were optimised based on trial-and-error on a smaller representative batch. The parameter values used for both segmentations are shown in Table 2.

**Table 2:** Segmentation method specific parameters used to analyse the synthetic data.

| Method |  |  |  |  |
| --- | --- | --- | --- | --- |
| pwvd | Window size (samples) | FM rate threshold (kHz/ms) | Acceleration threshold (kHz/ms<sup>2</sup>) | Extrapolation window(s) |
|  | 125 | 2 | 10 | 75x10e-6 |
| peak_percentage |  | peak percentage | double pass |  |
|  | 125 | 0.99 | True |  |

The accuracy of segmentation was determined by comparing the duration of the obtained call components and the original values used to make the synthesied calls. The accuracy of other parameters eg. CF peak frequency, FM bandwidth was not assessed. It follows directly that if the call components have been poorly segmented, any measurements made from the underlying audio will also be unrepresentative of the actual call parameters. Some calls appeared to have more than three components due to false positive CF/FM identifications, and were not included in the accuracy calculations.

### 0.3.2 Results

The `pwvd` method correctly identified 99% of all calls (322/324) as having only 3 components. The `peak_percentage` method correctly identified 94% of all calls as having 3 components (306/324). Both segmentation methods achieved a satisfactory performance. The `pwvd` method was superior in its segmentation accuracy to the `peak_percentage` method across all the parameter combinations and call components tested (Table 4, 3).

**Table 3:** Summary statistics describing the performance of the two segmentation methods on the synthetised test data set. The pwvd method performs better than the peak-percentage method over the tested parameter space and for all call components (iFM,tFM and CF).

| Call component | Region.type | Segmentation method | Segmentation accuracy: 95%ile range |
| --- | --- | --- | --- |
| CF | cf_duration | peak_percentage | [0.8994 1.05945] |
| CF | cf_duration | pwvd | [0.9392 1.02185] |
| tFM | downfm_duration | peak_percentage | [0.648 1.348] |
| tFM | downfm_duration | pwvd | [0.88675 1.208 ] |
| iFM | upfm_duration | peak_percentage | [0.624 1.3 ] |
| iFM | upfm_duration | pwvd | [0.918 1.2416] |

The relatively lower overall performance of the `peak_percentage` method can be specifically attributed to the call properties of certain synthetic calls. A further inspection of calls with lower than 0.8 accuracy in component duration revealed that calls with a high CF frequency (60 and 90 kHz) and at least one low bandwidth FM component (5kHz) were segmented with lower accuracy. This is explained by the fact that the peak-percentage of the recursive filter is set at 0.99 of the peak frequency. A low FM bandwidth call with a high CF frequency will have its cutoff frequency much below the actual CF frequency (600 and 900 Hz below peak frequency here). The lower cutoff frequency will thus lead to a shorter duration estimate of the low-bandwidth FM component. The accuracy of component durations was above 0.8 for all calls segmented with the `pwvd` method.

One percent of all `pwvd` segmented calls and six percent of all `peak_percentage` segmented calls had more than three detected call components. What caused the false positive call component detections in the `pwvd` and `peak_percentage` methods? In the `peak_percentage` method, the false component detections consisted of very short (≤0.1ms) falsely detected CF and FM segments located next to one another. These neighbouring CF and FM segments were caused by brief alterations in the dB rms levels of the high and low-passed audio. The brief alterations in the dB rms levels are likely due to the combination of windowing function applied on the synthetic calls and edge effects during high/low pass filtering. Such edge effects may not necessarily occur during the processing of experimentally recorded calls, which may have smoother roll-offs in call level. The two cases where false components were detected with the `pwvd` method were borderline cases where false CF components were detected in what should have been an FM region of the call. On further inspection it was shown that the frequency tracking of these false CF components was indeed accurate, but the action of the error-correction routines caused a slight drop in the frequency modulation rate to 1.9 kHz/ms, just slightly below the threshold of 2.0 kHz/ms. The error-correction routines in `pwvd` are typically required when low signal-level at the beginning and ends of the call causes jumps in the frequency tracking.

**Figure 4:**
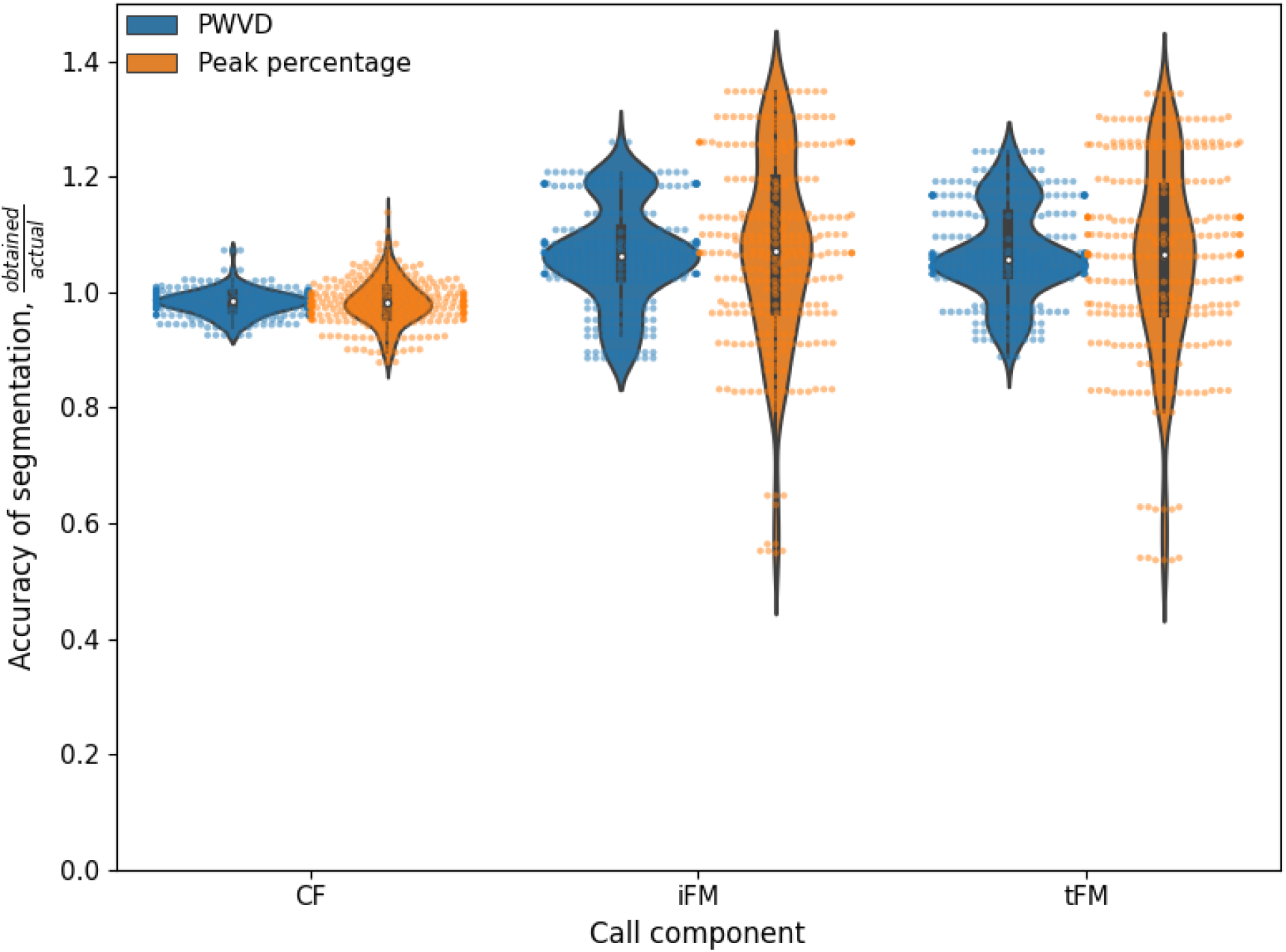
Accuracy of call component segmentation of the synthetic test data set shown with raw data overlaid on violinplots. The accuracy is calculated as the measured call component duration by the original duration. Blue violinplots: accuracy of the `pwvd` method, orange violinplots: accuracy of the peak-percentage method. The `pwvd` method is superior to the peak-percentage method in its segmentation performance across call components.

## 0.4 Discussion

Software based automation in acoustic analysis is an important step in ensuring reproducible results, which in turns spurs the growth of the research field (McFee et al. 2018; Baker and Vincent 2019). The `itsfm` package written in the Python (Van Rossum and Drake Jr 1995) language is an open-sourced method which may be used in the analysis of animal vocalisations such as CF-FM bat calls, and other vocalisations. The `itsfm` package has already been successfully used to segment and measure call parameters in an upcoming publication on group echolocation in CF-FM bats (Mysuru Rajagopalachari et al. 2020). The package introduces a new method the ‘pwvd’ method to segment CF and FM components based directly on the rate of frequency modulation. The ‘pwvd’ method also performs consistently better than the ‘peak-percentage’ method, and is thus the recommended segmentation method to use, at least for sounds that resemble CF-FM calls.

The use of `itsfm` in the analysis of other types of vocalisations still needs further explored. For instance, bird calls have been analysed (See online user-guide). The current ‘pwvd’ frequency tracking implementation only tracks a single frequency per point of time, and thus is not able to handle multi-harmonic sounds with equal harmonic emphasis very well. Future implementations of frequency tracking need to apply more sophisticated problem-region detection and also frequency tracking (eg. Viterbi path).

## 0.5 Open-source software and packages used

`itsfm` is written in the Python language (Van Rossum and Drake Jr 1995), and relies on the numpy, scipy, pandas, matplotlib and tftb (Oliphant 2006; Virtanen et al. 2020; Hunter 2007; McKinney and others 2010; Deshpande 2019). The Jupyter Notebook and Rmarkdown projects (Kluyver et al. 2016; Xie, Allaire, and Grolemund 2018) were used in the analysis of data and writing of this paper.

## 0.6 Supporting information

The `itsfm` package can be installed from the Python package index (PyPi) with the command `pip install itsfm`. The latest versions of the package and drafts of this paper are accessible at https://github.com/ thejasvibr/`itsfm`. Online documentation with detailed examples and troubleshooting guides can be accessed at https://itsfm.readthedocs.io

## 0.7 Acknowledgements

I would like to thank Diana Schoeppler for sharing know-how on analysing CF-FM calls and Neetash MR for helpful discussions. This work was funded by the DAAD and the IMPRS for Organismal Biology. I’d like to thank the following people for contributing to the call recording library Aditya Krishna, Aiqing Lin, Gloria Gessinger, Klaus-Gerhard Heller, Laura Stidsholt and Neetash MR.

